# Cell-associated, Heparin-like Molecules Modulate the Ability of LDL to Regulate PCSK9 Uptake

**DOI:** 10.1101/329722

**Authors:** Adri M. Galvan, John S. Chorba

**Author notes:** Contact information for corresponding author John S. Chorba 1001 Potrero Ave, Building 100, Room 261, Division of Cardiology, Zuckerberg San Francisco General, San Francisco CA 94110.

## Abstract

Proprotein convertase subtilisin/kexin type 9 (PCSK9) targets the LDL receptor (LDLR) for degradation, increasing plasma LDL and, consequently, cardiovascular risk. Uptake of secreted PCSK9 is required for its predominant effect on the LDLR. LDL itself inhibits this uptake, though the mechanism by which it does so remains unclear. In this study, we investigated the relationship between LDL, the PCSK9:LDLR interaction, and PCSK9 uptake. We show that LDL inhibits binding of PCSK9 to the epidermal growth factor precursor homology domain A (EGF-A) domain of the LDLR *in vitro* more impressively than it inhibits PCSK9 uptake in cells. Furthermore, a cell-based factor responsive to heparin-targeting treatments can explain this difference, consistent with its identity as a cell surface heparan sulfate proteoglycan (HSPG), a known co-receptor for PCSK9. Furthermore, we show that the entire PCSK9 prodomain, but not truncated variants, rescues PCSK9 uptake in the presence of LDL, suggesting that PCSK9:LDL binding requires the entire prodomain. Additionally, we show that the gain-of-function (GOF) PCSK9 variant S127R has increased affinity for heparin-like molecules such as HSPGs, potentially explaining the biochemical basis for its phenotype. Overall, our findings suggest a model where PCSK9, LDL, and HSPGs all interact to regulate PCSK9 uptake into the hepatocyte.

## Introduction

Gain-of-function (GOF) mutations in the proprotein convertase subtilisin/kexin type 9 (PCSK9) gene, which encodes a self-cleaving protease, cause autosomal dominant familial hypercholesterolemia (1). Mechanistically, PCSK9 binds the epidermal growth factor precursor homology domain A (EGF-A) of the LDL receptor (LDLR) (2) and induces lysosomal-mediated degradation of the complex (3), thus limiting the availability of LDLR to recycle back to the cell surface and internalize LDL (4). The robust relationship between LDL and atherosclerosis (5) combined with a lack of ill effects in individuals devoid of functional PCSK9 (6, 7) has made PCSK9 an important therapeutic target against heart disease. Indeed, monoclonal antibodies inhibiting PCSK9 reduce serum LDL levels (8, 9), and also reduce cardiovascular events (10), even when added to current effective therapies such as statins. Despite this impressive efficacy, these biologic therapies suffer from a lack of cost-effectiveness and a requirement for subcutaneous injections, leaving a need for additional therapeutic options (11, 12). Indeed, other strategies to inhibit PCSK9 function are under investigation (13). Since atherosclerosis is a chronic disease, a successful therapy may potentially be given for many years.

Despite the clear effects of PCSK9 on the LDLR and serum LDL, certain aspects of PCSK9 biology remain incompletely understood (14). Understanding this biology may uncover new pathways amenable to certain therapeutic strategies, or presage long-term effects of therapies which we might not otherwise foresee. Why PCSK9 exists, for example, remains an unanswered question (15), as it seems rather odd for the predominant function of a gene to cause disease. In this vein, a series of unrelated studies have shown LDL to bind PCSK9 both intra and extracellularly, augmenting secretion of apoB containing lipids, (16) and inhibit the ability of serum PCSK9 to induce LDLR degradation in humans (17). Intriguingly, the N-terminus of the PCSK9 prodomain, which is dispensable for the PCSK9:LDLR interaction (18, 19), is required for PCSK9 to bind LDL (17). One model proposes that PCSK9 inhibits the interaction between apoB containing particles and the LDLR in the secretory pathway so as to allow both components to reach the extracellular space (16).

Regulation of PCSK9 occurs at both the transcriptional (20–23) and post-transcriptional levels (14), with secretion of PCSK9 itself an important regulatory step for its function (24). As a proprotein convertase, PCSK9 undergoes self-proteolysis as a requirement to permit exit from the endoplasmic reticulum (ER) (25–27), serving as the overall rate-limiting step of PCSK9 secretion (28). Protease-dead PCSK9 mutants do not undergo efficient secretion and have minimal effect on the LDLR (29). Despite the general requirement for self-cleavage and secretion for full PCSK9 function, the initial GOF PCSK9 mutant identified, S127R, has long been known to have paradoxically reduced self-processing, suggesting additional mechanisms can overcome this functional restriction (26, 30). Recently, heparan sulfate proteoglycans (HSPGs) have been identified as co-receptors for PCSK9 on the hepatocyte cell surface, with a specific arginine-rich motif encompassing residues 93-139 in the PCSK9 prodomain as the likely binding site mediating this interaction (31).

Given the involvement of the PCSK9 prodomain in both interactions, we postulated that potentially overlapping binding sites for LDL and HSPGs could serve a regulatory role for PCSK9 function. In this study, we investigated the relationship between LDL as an inhibitor of PCSK9 uptake into hepatocytes. We found that while LDL is a potent inhibitor of PCSK9 binding to the LDLR, its ability to inhibit PCSK9 uptake into cells is less impressive. We provide evidence that this reduction in inhibitory potential is related to cell-based heparin-like molecules, and that the region of PCSK9 required for LDL to inhibit its uptake overlaps with PCSK9’s binding site for HSPGs. Furthermore, we show that the S127R mutant of PCSK9 shows increased affinity for heparin, suggesting that increased affinity for HSPGs may explain its GOF phenotype. Lastly, we show that differences in LDL variant density do not affect PCSK9 uptake. Overall, our findings provide support to a model wherein PCSK9, LDL, and HSPGs interact in a complex manner to modulate lipid homeostasis in the hepatocyte.

## Materials and Methods

### Plasmid Construction

All expression vectors were created by Gibson assembly (32) after PCR amplification of appropriate PCSK9 domains from previously described plasmids (27, 28), a plasmid encoding nanoluciferase (NLuc) (Promega, Madison WI), and a plasmid encoding the Fc-Avi tag (genereous gift from A. Martinko and J. Wells, UCSF). The PCSK9-NLuc plasmid and its variants were placed into the pcDNA5/FRT/TO backbone (ThermoFisher Scientific, Waltham MA), and the ProPCSK9-Fc-Avi plasmid and its variants were placed into the pcDNA3.4 backbone (ThermoFisher Scientific). Insertion of mutations (S127R, D374Y) and truncations was performed by site-directed mutagenesis (33) using custom synthesized oligonucleotide primers (Elim Biopharmaceuticals, Hayward CA). All constructs were extensively sequenced to ensure the absence of errors.

### Cell Culture

HEK293T cells (ATCC, Manassas VA) were maintained in high-glucose, pyruvate, and L-glutamine containing DMEM (ThermoFisher Scientific) supplemented with 10% FBS (Axenia BioLogix, Dixon CA) at 37 °C under 5% CO_2_ and dissociated for passage by 0.05% Trypsin-EDTA (ThermoFisher Scientific). HepG2 cells (ATCC) were grown in low-glucose, pyruvate, and GlutaMAX containing DMEM (ThermoFisher Scientific) supplemented with 10% FBS at 37 °C under 5% CO_2_. To minimize cell clumping, HepG2 cells were dissociated by 0.25% Trypsin-EDTA (ThermoFisher Scientific) and sent three times through a 21 gauge needle during each passage. For several experiments requiring upregulation of cell-surface LDL receptors, HepG2 cells were treated with sterol-depleting medium, which consisted of low-glucose, pyruvate, and GlutaMAX containing DMEM supplemented with 5% lipoprotein-deficient serum (Kalen BioMedical, Germantown MD), 25 µM mevastatin (MilliporeSigma) and 50 µM mevalonolactone (MilliporeSigma). Flp-In T-Rex 293 cells (ThermoFisher Scientific) were maintained in the same conditions as 293T cells, but additionally supplemented with 1 mg/ml Zeocin (InvivoGen, San Diego CA) and 15 µg/ml blasticidin (InvivoGen) prior to selection, or 150 µg/ml Hygromycin B (InvivoGen) and 15 µg/ml blasticidin after selection of stable cell lines.

### PCSK9-NLuc and ProPCSK9-Fc-Avi Media Production

HEK293T cells were seeded at 1 × 10^6^ cells per T25 flask and incubated overnight. On the following day, the cells were transfected with Lipofectamine 3000 (ThermoFisher Scientific) according to the manufacturer’s protocol using 6944 ng of appropriate plasmid DNA per well. The medium was changed 6–12 h after transfections and harvested another 24 h later. Relative amounts of PCSK9-NLuc were quantitated by luminescence assay (see below). Relative amounts of ProPCSK9-Fc-AVI were quantitated using a Human Fc ELISA Kit (Syd Labs, Natick MA) according to the manufacturer’s instructions. For some experiments, a stable, inducible PCSK9-NLuc cell line, derived from the Flp-In T-Rex 293 cell line (ThermoFisher Scientific) according to the manufacturer’s instructions, was induced with 1 µg/ml doxycycline (MilliporeSigma, Burlington MA) and conditioned medium was collected 24-48 h later.

### Luminescence Assays

Luciferase containing samples in a white 96 well plate were mixed with 60 µl of a filtered 2× coelenterazine reagent for readout (300 mM sodium ascorbate (MilliporeSigma), 5 mM NaCl, 40 µM coelenterazine (Gold Biotechnology, St. Louis MO), and either 0.1% BSA (MilliporeSigma) for media-based non-lytic assays or 0.1% Triton X-100 (MilliporeSigma) for cell-based lytic assays. The samples were incubated at room temperature in the absence of light with gentle shaking shaker for 5 to 10 min followed by immediate readout of luminescence on a plate reader (Tecan Systems, San Jose CA).

### PCSK9 in vitro Binding Assays

Inhibition of the PCSK9-LDLR interaction by LDL (Lee BioSolutions, Maryland Heights MO) was evaluated using a CircuLex PCSK9-LDLR *in vitro* binding assay kit (MBL International, Woburn MA) according to the manufacturer’s instructions with minor modifications. PCSK9 was incubated with LDL for 1 h prior to use in the assay. Relative PCSK9:LDLR(EGF-A) binding was defined as absorbance at 450 nm, with a value of 1 defined as the highest concentration of PCSK9 in the absence of LDL. To evaluate PCSK9 binding to heparin, conditioned media containing PCSK9-NLuc and its variants were incubated in clear heparin-coated microplates (Bioworld, Dublin OH) at room temperature with gentle shaking for 2 h. Input luminescence was measured as described above. Each well was then washed 2× with 500 mM NaCl, and luminescence assay was performed. The wash protocol was repeated with 5 mg/ml heparin (MilliporeSigma) and luminescence assay was repeated.

### PCSK9-NLuc Uptake Assay

HepG2 cells were seeded into 12-well plates at 5 × 10^5^ cells per well with 0.5 ml of sterol-depletion medium and incubated overnight. Heparinase-treated cells were incubated with heparinase (MilliporeSigma) at 0.1-0.5 U/ml for 1 h and washed with PBS. All cells were provided fresh medium on the day of experiment, supplemented with 50 µM chloroquine (MilliporeSigma) as appropriate to inhibit lysosome-mediated PCSK9 degradation, and then treated with appropriate conditioned PCSK9-NLuc media and/or LDL treatments as indicated. Conditioned PCSK9-NLuc media was normalized by luminescence assay prior to co-incubation to ensure uniform treatment for each experiment. Typically, treated cells were incubated with treatments at 37 °C for 4 h, washed, and dissociated with 0.25% Trypsin-EDTA. The cells were centrifuged at 300 × *g* for 5 min to remove supernatant, and lysed in in lysis buffer (50 mM Tris-HCl pH 7.4, 150 mM NaCl, 1× cOmplete protease inhibitor (MilliporeSigma), 0.1% Nonidet P-40), and then clarified from insoluble fraction at 21,000 × *g* for 15 min. Protein concentration was analyzed by a Micro BCA Protein Assay Kit (ThermoFisher Scientific), and equal amounts of lysate were then evaluated by luciferase assay. For experiments utilizing methylated LDL, LDL was reductively methylated as described in the literature.(34, 35) Briefly, LDL (10mg/ml) in 0.15 M NaCl was treated with 1.5 volumes of 0.3 M sodium borohydride followed by six additions of 1 µl of 37% aqueous formaldehyde over 30 min at 4 °C. Small dense and large LDL were separated from LDL using a one-layer system with ultracentrifugation (350,000 × *g*) over a OptiPrep density gradient medium (CosmoBio, Tokyo Japan) as per prior literature protocols.(36)

### LDL Uptake Assays

HepG2 cells were seeded at 1 × 10^5^ cells per well in a 12 well plate in 0.5 ml sterol-depletion medium and incubated overnight. The medium was then replaced with fresh sterol-depletion medium or 0.5 mL of conditioned medium containing PCSK9-NLuc D374Y and incubated for 4 h. 10 µg/ml BODIPY FL LDL (ThermoFisher Scientific) was added to each well and incubated for 2 h. The cells were washed three times with PBS, dissociated with 0.25% Trypsin-EDTA, collected, centrifuged, lysed, and clarified as in the PCSK9 uptake assay. Equal concentrations of lysates were added to a black clear bottom 96 well plate and fluorescence (λ_em_ = 515 nm) was analyzed on a plate reader.

### Data Analysis

Data and statistical analysis was performed using Prism 7.0 (GraphPad Software, La Jolla CA). In general, raw luminescence or fluorescence outputs were normalized to positive and negative control samples, and experiments were performed in triplicate and repeated at least three times. Error bars in all graphs represent standard deviation (SD).

## Results

### LDL inhibits the in vitro binding of PCSK9 to the LDLR more potently than PCSK9 uptake in tissue culture

We first sought to evaluate the inhibitory effect of LDL on complex formation between PCSK9 and the EGF-A domain of the LDLR. Prior gel shift studies from others had shown that LDL did not disrupt the PCSK9:LDLR(EGF-A) interaction (17), though these studies were conducted at supra-physiologic levels of PCSK9, using a GOF D374Y variant with increased affinity for the EGF-A domain (18). We thus employed a commercially available ELISA using near physiologic levels of PCSK9 and saturating levels of EGF-A domain to probe the interaction (37). In contrast to prior results, we found that both sub-physiologic and physiologic levels of LDL potently inhibited PCSK9:LDLR(EGF-A) binding in a dose-dependent manner (Fig. 1A). Specifically, we found that at PCSK9 concentrations of 1.4 nM (100 ng/ml), 10 mg/dl of LDL inhibited PCSK9:LDLR(EGF-A) domain binding by 75% (Fig. 1A, light blue) while 100 mg/dl of LDL inhibited binding by 93% (Fig. 1A, dark blue).

**Figure 1:**
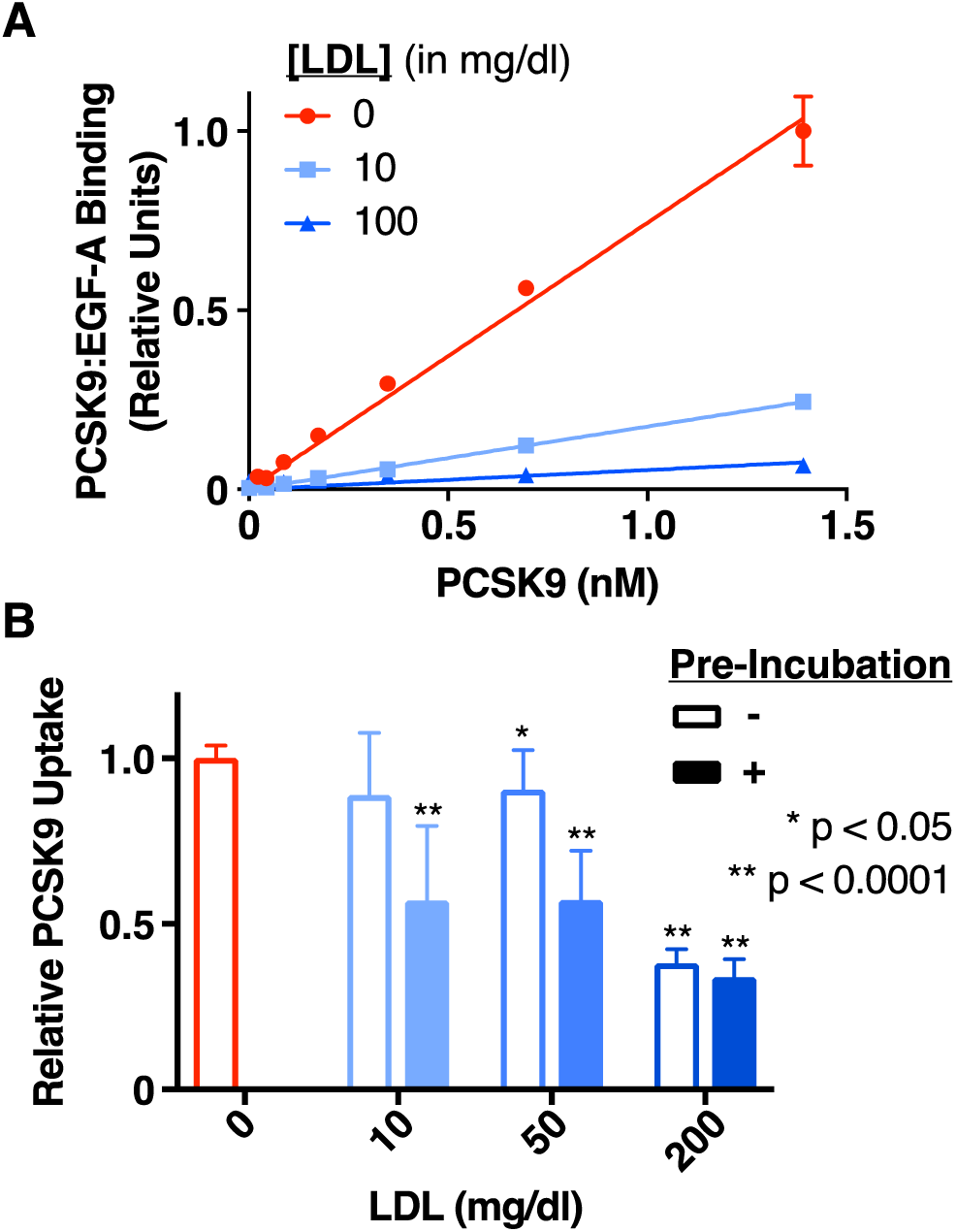
Effect of LDL on the PCSK9:LDLR Interaction and PCSK9 Uptake. A, Relative binding of increasing concentrations of PCSK9 to the EGF-A domain of LDLR in the absence (red) or presence of sub-physiologic (10 mg/dl, light blue) or physiologic (100 mg/dl, dark blue) LDL, as measured by a commercially available PCSK9-LDLR ELISA assay. Raw absorbance data was normalized to 1 for the highest PCSK9 concentration in the absence of LDL and 0 in the absence of PCSK9. The linear regression lines shown were generated using Prism 7 software, with the extra-sum-of-squares F test indicating different slopes at p < 0.0001. B, Relative uptake, as measured by luminescence assay, of PCSK9-NLuc by HepG2 cells in the presence of increasing concentrations of LDL, shown by sequential additions of PCSK9 and LDL to cells (open bars) or by pre-incubation of PCSK9 with LDL prior to addition to cells (filled bars). Raw luminescence data was normalized to 1 for PCSK9-NLuc treatment in the absence of LDL and 0 in the absence of PCSK9-NLuc uptake. P values indicate results of an unpaired t-test with Welch’s correction compared to the control (no LDL) arm.

As PCSK9 uptake is dependent on the binding to cell surface LDLRs (38), we then sought to extend our *in vitro* findings to a cellular setting. To focus specifically on PCSK9 binding and uptake, we engineered a luciferase-tagged PCSK9 (PCSK9-NLuc) to add *in trans* to HepG2 cells, which were treated with chloroquine to inhibit lysosome-mediated PCSK9 degradation (39). The luciferase-based system allowed us to control the relative input from the conditioned media of 293T cells overexpressing the tagged PCSK9, and ultimately measure the luciferase activity of the treated HepG2 lysates as a proxy for PCSK9 uptake. To ensure our system had the desired downstream physiologic effect on the LDLR, we confirmed that HepG2 cells treated with PCSK9-NLuc D374Y reduced LDL uptake compared with untreated controls, using a fluorescence based BODIPY LDL uptake assay (Fig. S1).

We then employed our system to evaluate the effects of LDL on PCSK9 uptake. We incubated conditioned PCSK9-NLuc medium with chloroquine-treated HepG2 cells in the presence of sub-physiologic (10 mg/dl), low (50 mg/dl), and high (200 mg/dl) concentrations of LDL. In contrast with the results of the *in vitro* ELISA assay, LDL inhibited PCSK9 uptake modestly, with a marked effect only at the high concentration (Fig. 1B, open bars). We repeated our experiments by pre-incubating the conditioned PCSK9-NLuc media with LDL prior to adding the mixture to HepG2 cells. This resulted in much more impressive reductions of PCSK9 uptake (Fig. 1B, filled bars), partially rescuing the inhibitory activity of LDL, particularly for the 10 mg/dl and 50 mg/dl concentrations, though not quite to the levels seen in the ELISA alone. We conclude from this data that a cell-based factor partially abrogates the ability of LDL to reduce PCSK9 uptake into cells. Furthermore, this cell-based factor appears most effective when PCSK9 and LDL are presented simultaneously to the cells.

### LDL mediated inhibition of PCSK9 uptake is sensitive to reagents which modify the interaction between LDL and heparin-like molecules

Our initial results suggested that the cell-based factor might be directly competing with LDL to bind PCSK9 on a relatively slow kinetic timescale. We hypothesized that HSPGs on the HepG2 cells may be this cell-based factor, given the recent description of their affinity for the PCSK9 prodomain and involvement in PCSK9 uptake (31). To test this, we evaluated PCSK9 uptake with reagents expected to modify HSPG activity. First, we treated HepG2 cells with heparinase to remove cell surface heparin-like molecules (including HSPGs) prior to evaluating the effect of LDL on our PCSK9-NLuc uptake assay. Unsurprisingly, and consistent with prior results (31), heparinase treatment in the absence of LDL reduced PCSK9-NLuc uptake (Fig. 2A, orange), with its absolute effect similar to that of pre-incubation of PCSK9 with LDL. Physiologic levels of LDL added to heparinase treated cells reduced PCSK9 uptake further (Fig. 2A, brown), suggesting that at least to some extent, LDL can inhibit PCSK9 uptake even in the absence of HSPGs.

**Figure 2:**
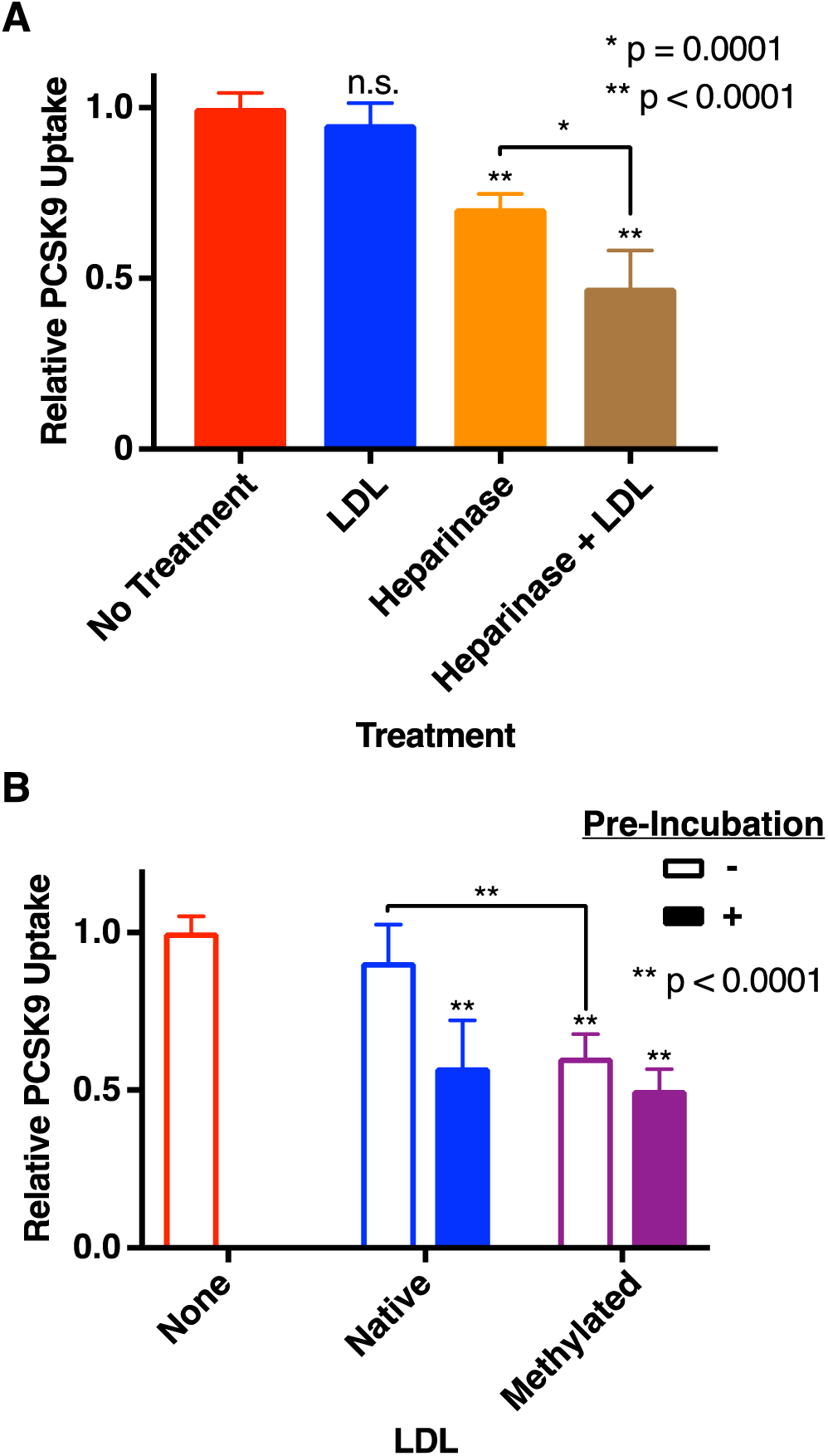
Sensitivity of PCSK9 Uptake to Heparinase and Reductively Methylated LDL. A, Relative uptake, as measured by luminescence assay, of PCSK9-NLuc by HepG2 cells co-incubated with LDL (blue), treated with heparinase (orange), or both (brown). Raw luminescence data was normalized to 1 for PCSK9-NLuc treatment in the absence of LDL or heparinase treatment and 0 in the absence of PCSK9-NLuc. P values indicate results of an unpaired t-test with Welch’s correction, compared to the control arm (no treatment, red) unless otherwise indicated. B, Relative uptake, as measured by luminescence assay, of PCSK9-NLuc by HepG2 cells in the presence of either native (blue) or reductively methylated (purple) LDL, stratified by sequential addition (open bars) or pre-incubation (filled bars) of PCSK9 with LDL pre-incubation. Raw luminescence data was normalized to 1 for PCSK9-NLuc treatment in the absence of LDL and 0 in the absence of PCSK9-NLuc uptake. P values indicate results of an unpaired t-test with Welch’ correction, compared to the control arm (no LDL, red) unless otherwise indicated.

As native LDL binds negatively charged sulfated glycoproteins (such as HSPGs) (34, 40, 41), we also hypothesized that competition between cellular HSPGs and PCSK9 for LDL binding could explain our differential effects on pre-incubation. We thus subjected LDL to reductive methylation, which reduces the affinity of LDL to bind either HSPGs or the LDLR itself (34, 35). Consistent with our hypothesis, methylated LDL retained the ability to inhibit PCSK9 uptake to levels similar to pre-incubation with native LDL (Fig. 2B). Overall, when taken together with data from heparinase treated cells, these data suggest that cell-based heparin-like molecules, such as HSPGs, compete with LDL to bind PCSK9.

### The full-length PCSK9 prodomain can outcompete the inhibitory effect of LDL on PCSK9 uptake

Prior work has shown the N-terminus of the prodomain, specifically residues 31-52, as necessary for LDL binding (17). Deletion of these residues also improves PCSK9 binding to soluble LDLR (18), and increases the ability of PCSK9 to downregulate LDLR on cells (19). By contrast, the binding site of HSPGs to the PCSK9 prodomain encompasses an arginine-rich motif spanning residues 93 to 139 (31). To ask whether the N-terminal prodomain residues were sufficient to mediate the inhibitory effect of LDL on PCSK9 uptake (and therefore non-overlapping with the HSPG binding site), we created secreted PCSK9 prodomains to use in competition experiments. We designed our prodomain constructs as N-terminal fusions to secretable human Fc domains, a strategy similar to one previously used to permit secretion of the prodomain in the absence of the PCSK9 catalytic domain (42). Using the previously described binding sites as our guide, we generated three sequential C-terminal truncations of the prodomain, encompassing residues 31-50, 31-100 and 31-152 (Fig. 3A). We expressed these prodomain-Fc fusions via transient transfections in 293T cells and evaluated their effects on LDL mediated PCSK9 uptake inhibition. Our results showed that complete rescue of PCSK9 uptake required the full (i.e. non-truncated) prodomain (Fig. 3B). These data are consistent with the requirement of the entire prodomain, including the HSPG binding site, to permit the LDL particle to inhibit PCSK9 uptake.

**Figure 3:**
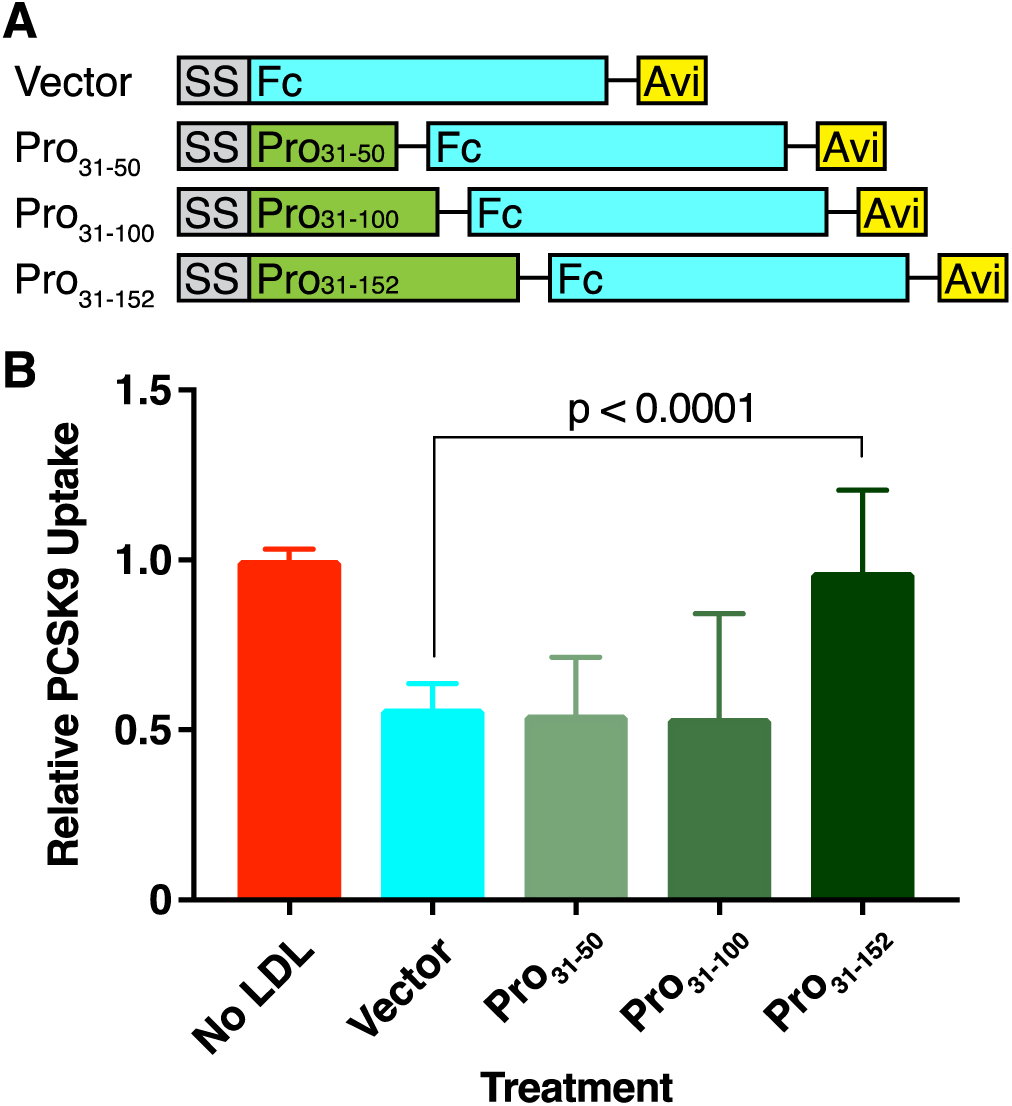
Rescue of LDL-Mediated Inhibition of PCSK9 Uptake by Exogenous Prodomain. A, Schematic of exogenous prodomain constructs. All constructs are driven by a CMV promoter (not shown), and include the PCSK9 signal sequence (SS, grey) and C-terminal Fc (cyan) and Avi (yellow) tags. The PCSK9 prodomain variants (green) include residues 31-50, 31-100, or 31-152 (full-length). B, Relative uptake, as measured by luminescence assay, of PCSK9-NLuc by HepG2 cells in the presence of LDL and the indicated exogenous prodomain Fc-Avi fusion. Note that in these experiments, all LDL was pre-incubated with the PCSK9-NLuc and exogenous prodomain. Raw luminescence data was normalized to 1 for PCSK9-NLuc treatment in the absence of LDL and 0 in the absence of PCSK9-NLuc uptake. P values indicate the results of an unpaired t-test with Welch’s correction.

### The gain of function S127R PCSK9 variant has increased affinity for heparin and causes increased PCSK9 uptake

The initial description of PCSK9 as a Mendelian cause of familial hypercholesterolemia came from the PCSK9 S127R mutant (1). Biochemical characterization of this mutant has remained puzzling; both self-proteolysis and secretion are required for PCSK9 to exert its maximal effect on the LDLR, yet we and others have shown that the S127R variant is deficient in both proteolytic function and secretion (28, 43). Given the location of S127 within the prodomain and HSPG binding motif, we hypothesized that a positively charged arginine at this position could increase the affinity of PCSK9 for the negatively charged HSPGs, potentially explaining the GOF phenotype. To address this, we first evaluated the affinity of PCSK9 S127R for heparin in an *in vitro* assay. We generated WT, S127R, and D374Y PCSK9-NLuc constructs, and produced the corresponding proteins via transient transfections in 293T cells. We then added the conditioned media to a heparin-coated plate and measured the relative luminescence of the PCSK9 inputs. We sequentially washed the plates with a high salt solution followed by a competitive, free heparin solution, reading out luminescence after each step to monitor the remaining bound PCSK9-NLuc. Consistent with our hypothesis, a 4-fold greater proportion of S127R PCSK9-NLuc, compared to the WT and D374Y variant, remained bound to the heparin plate after the salt wash (Fig. 4A). Furthermore, the S127R variant approached the basal level after the competitive wash with excess free heparin (Fig. 4A).

**Figure 4:**
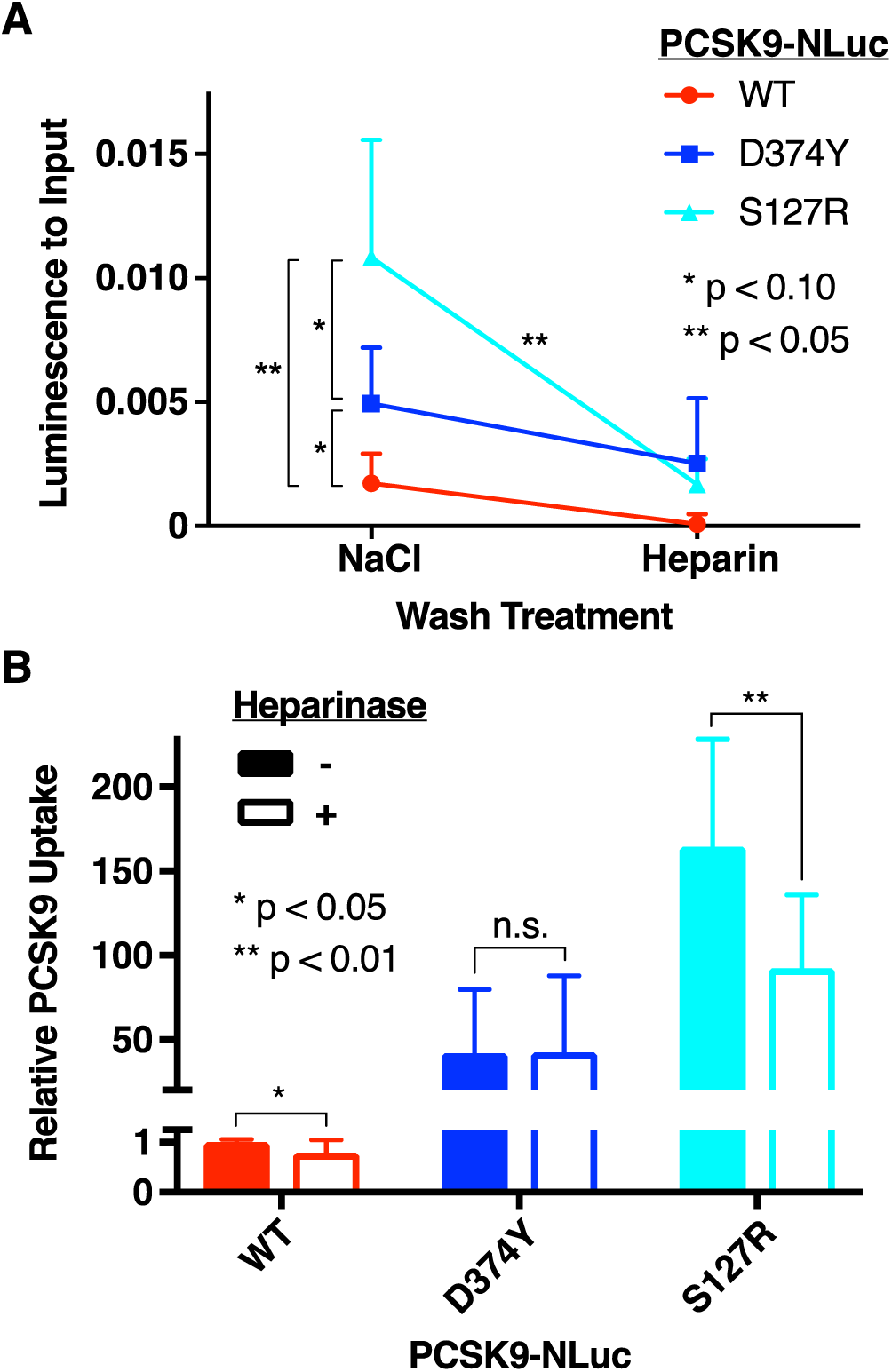
Effect of PCSK9 S127R on Heparin Binding and PCSK9 Uptake. A, Relative *in vitro* binding affinity of WT (red), D374Y (blue), and S127R (cyan) PCSK9-NLuc to heparin after high-salt and heparin competition washes, as measured by luminescence assay. The Y axis shows relative luminescence compared to initial input on the heparin-coated plate. P values indicate the results of an unpaired t-test with Welch’s correction. Only one side of each set of error bars is shown for clarity. B, Relative PCSK9 uptake, as measured by luminescence assay, by heparinase untreated (filled bars) and treated (open bars) HepG2 cells incubated with WT (red), D374Y (blue) or S127R (cyan) PCSK9-NLuc conditioned media. Luminescence data was normalized to 1 for WT PCSK9-NLuc treatment 0 in the absence of PCSK9-NLuc treatment. Note the discontinuous Y axis, which highlights the increased uptake of the PCSK9 variants compared to WT. P values indicate the results of an unpaired t-test with Welch’s correction.

We then asked whether these findings would translate to cellular uptake of PCSK9. To do so, we repeated our cellular uptake assays using the WT, S127R, and D374Y PCSK9-NLuc variants with heparinase-treated and untreated HepG2 cells. Our results show that S127R and D374Y PCSK9 mutants show increased uptake in the absence of heparinase treatment, consistent with the known increased affinity of D374Y PCSK9 for the LDLR (Fig. 4B, filled bars). Moreover, the S127R mutant is more sensitive to heparinase treatment than either WT or the D374Y mutant (Fig. 4B, open bars). These results are consistent the mechanism of S127R uptake being dependent on a heparinase-sensitive factor such as cell surface HSPGs.

### LDL density variants do not differentially affect PCSK9 uptake

We next evaluated the impact of LDL density variants on the inhibitory mechanism of PCSK9 uptake. Size and density of LDL has been identified as a factor which can alter the atherogenicity of the LDL particle (44–47), and thus we investigated the hypothesis that differential effects on PCSK9 uptake could mediate this propensity towards atherosclerosis. We generated small dense LDL (sd-LDL) and large LDL (l-LDL) by ultracentrifugation, and compared the effects of each in both the PCSK9:LDLR(EGF-A) *in vitro* binding ELISA and cell-based PCSK9-NLuc uptake assays. Our results showed no significant differences in the inhibition of PCSK9:LDLR(EGF-A) binding *in vitro* (Fig. 5A), nor in PCSK9 uptake between either species (Fig. 5B).

**Figure 5:**
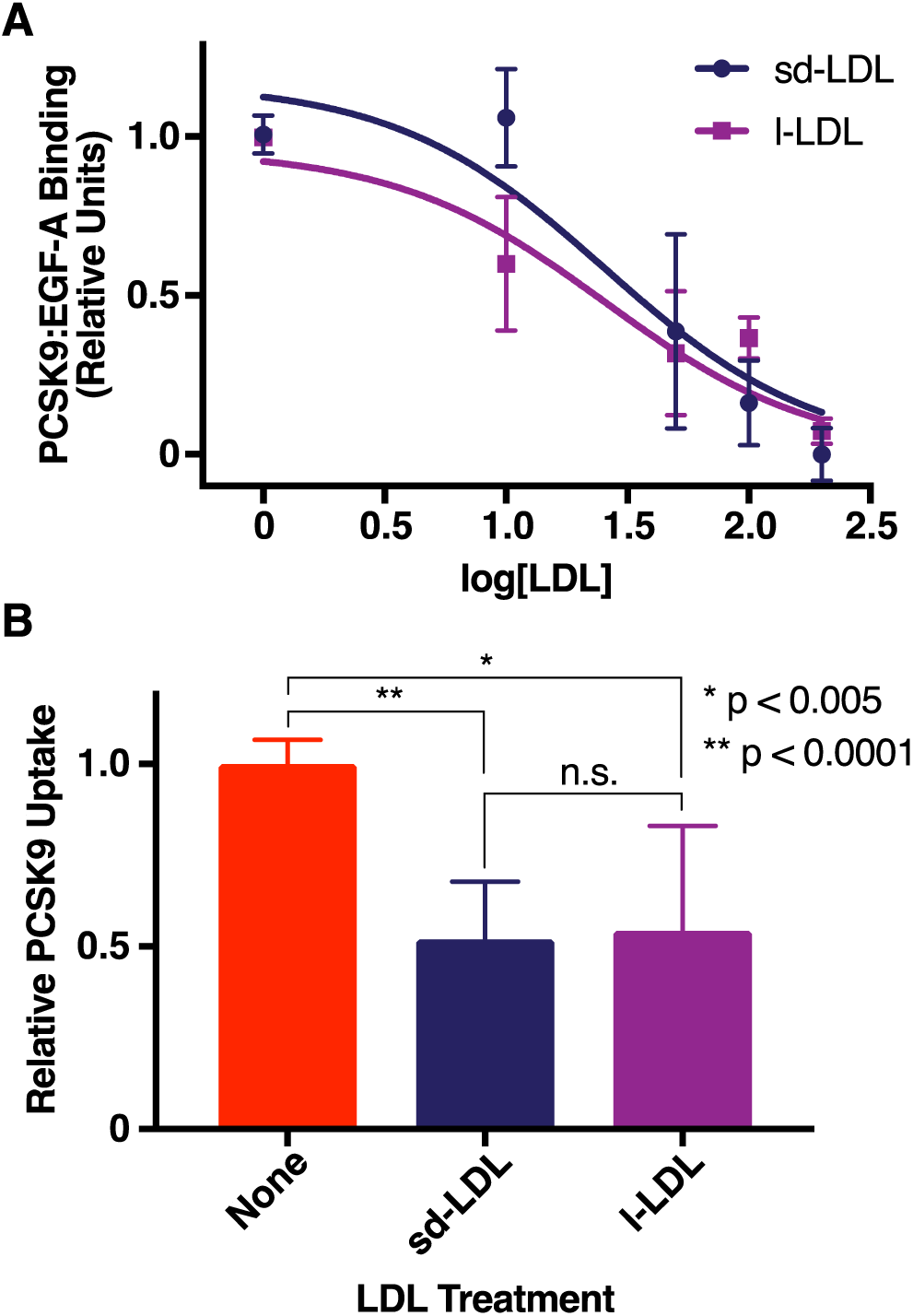
Effect of LDL Variants on PCSK9:LDLR Binding and PCSK9 Uptake. A, Relative binding of PCSK9 to the EGF-A domain of LDLR in increasing concentrations of sd-LDL (indigo) or l-LDL (purple), as measured by a commercially available PCSK9-LDLR ELISA assay. Raw absorbance data was normalized to 1 for the maximal absorbance and 0 for minimal absorbance. Nonlinear regression curves, constrained to a minimum of 0, were generated using Prism 7 software, with the extra-sum-of-squares F test failing to reject the hypothesis (p = 0.85) that one curve fits both data sets. B, Relative PCSK9 uptake, as measured by luminescence assay, of PCSK9-NLuc by HepG2 cells in the presence of sd-LDL (indigo) or l-LDL (purple). Note that PCSK9-NLuc was pre-incubated with the LDL variants prior to co-incubation with the cells. P values indicate the results of unpaired t-tests with Welch’s correction.

## Discussion

In this study, we examined the role of LDL as an inhibitor of PCSK9 uptake. Consistent with prior studies, we found that LDL indeed inhibits PCSK9 uptake in cells, but in contrast with prior studies, we found that LDL inhibits PCSK9:LDLR(EGF-A) binding *in vitro* (17). We suspect that the differences between our findings and those of previous studies are explained by differences in experimental detail. First, our study used over 100-fold lower concentrations of PCSK9, and thus a much higher ratio of LDL to PCSK9. Second, we utilized an ELISA approach, as opposed to a gel-shift assay. The wash steps in the ELISA would remove any free PCSK9, as well as free LDL, potentially altering the binding equilibrium compared with a one-pot incubation. Though our findings suggest that the PCSK9:LDL interaction reduces the affinity LDLR for PCSK9, our findings remain consistent with the presence of separate PCSK9 binding sites for LDL and LDLR (17). Indeed, the large size of the LDL particle could either sterically or electrostatically hinder LDLR binding despite directly binding a remote region, or could otherwise induce an allosteric change to the LDLR binding site.

Regardless of these experimental differences, we found that the inhibition of PCSK9:LDLR(EGF-A) binding *in vitro* was more impressive than the degree of inhibition of PCSK9 uptake in cells. Though PCSK9 uptake is dependent upon the PCSK9:LDLR interaction (38), these results imply that an alternative cell-based factor is playing a role, either in modulating PCSK9:LDLR binding or PCSK9 uptake directly. We found that inhibition of PCSK9 uptake by LDL was sensitive to pre-incubation of the LDL with PCSK9 (when LDL levels were not high), as well as heparinase treatment of the recipient cells. Furthermore, reductive methylation of LDL, which reduces its affinity for heparin-like molecules, also rescued LDL inhibition of PCSK9 uptake without the need for pre-incubation. While reductive methylation also reduces the affinity of LDL for the LDLR itself, abrogation of this interaction alone cannot account for the reduction in PCSK9 uptake, since uptake remains higher in cells not treated with additional LDL beyond that present in the sterol-depleting medium. When taken together, these findings are consistent with HSPGs acting as cell-based factors to modulate PCSK9 uptake in the presence of LDL. In addition, competitive experiments using exogenous PCSK9 prodomain truncations show that full-length prodomain, but not the truncated prodomains, can rescue PCSK9 uptake to full levels in the presence of LDL. These data are most consistent with a requirement of the entire prodomain for PCSK9:LDL binding.

Though speculative, it is tempting to propose a model to incorporate all of this data. One relatively parsimonious explanation involves the negatively charged N-terminus of the PCSK9 prodomain (and specifically residues 31-52) in a fluid binding equilibrium with the positively charged, arginine-rich HSPG binding motif of residues 93-139. Binding of the prodomain to LDL could directly inhibit HSPG binding itself. Furthermore, the prodomain:LDL interaction could induce an allosteric change in the catalytic domain, thereby remotely affecting PCSK9:LDLR binding, potentially explaining the effect of further reduced PCSK9 uptake by LDL in heparinase-treated cells. HSPGs themselves could also serve to sequester LDL particles from PCSK9 at physiologic concentrations, particularly if differential binding kinetics are at play. An off-rate of PCSK9:LDL slower than the on-rate of HSPG:LDL, for example, could explain the sensitivity of our cell-based assay to pre-incubation of LDL and PCSK9.

Additionally, we investigated the role of the S127R mutant, given that the biochemical mechanism to explain its GOF phenotype has remained elusive. We provide evidence that the S127R mutant shows increased affinity for heparin, which we suspect translates to increased affinity for cell-surface HSPGs. Despite the reduction in processing efficiency, the S127R mutant still affects LDLR levels after being secreted (30), and an increased affinity for the HSPG co-receptor would be a tantalizing explanation. Interestingly, a nearby mutant, D129G, shows a similar phenotype (43). While not tested directly in our study, the removal of a negatively charged aspartate from the same binding region could potentially increase affinity for negatively charged HSPGs.

Overall, the interactions between LDL, PCSK9 and HSPGs suggest a complex interplay to regulate ultimate PCSK9 function. Further understanding of this interplay, and these regulatory mechanisms, may allow the identification of potential targets which could ultimately upregulate the LDLR, lower serum LDL, and protect against atherosclerotic disease.

## Acknowledgements and grant support

We thank Kevan Shokat (University of California San Francisco) for guidance, generous support, and critical reading of the manuscript. We also thank Sampath Parthasarathy (University of Central Florida) for suggesting the methylation of LDL and members of the Shokat laboratory for helpful discussion. This work was also supported by the Howard Hughes Medical Institute Medical Research Fellows Program (to AMG), the NIH/NHLBI (K08 HL124068 and LRP HMOT1243, both to JSC), the Hellman Foundation (to JSC) and a Pfizer ASPIRE Cardiovascular Award (to JSC). This content is solely the responsibility of the authors and does not necessarily represent the official views of the funding agencies.

## Abbreviations

PCSK9: proprotein convertase subtilisin/kexin type 9
LDLR: low density lipoprotein receptor
EGF-A: epidermal growth factor precursor homology domain A
HSPG: heparan sulfate proteoglycan
GOF: gain-of-function
ER: endoplasmic reticulum
NLuc: nanoluciferase
SD: standard deviation
sd-LDL: small dense LDL
l-LDL: large LDL

## References

1. Abifadel, M., M. Varret, J.-P. Rabès, D. Allard, K. Ouguerram, M. Devillers, C. Cruaud, S. Benjannet, L. Wickham, D. Erlich, A. Derré, L. Villéger, M. Farnier, I. Beucler, E. Bruckert, J. Chambaz, B. Chanu, J.-M. Lecerf, G. Luc, P. Moulin, J. Weissenbach, A. Prat, M. Krempf, C. Junien, N. G. Seidah, and C. Boileau. 2003. Mutations in PCSK9 cause autosomal dominant hypercholesterolemia. Nat. Genet. 34: 154–6.

2. Zhang, D.-W., T. A. Lagace, R. Garuti, Z. Zhao, M. McDonald, J. D. Horton, J. C. Cohen, and H. H. Hobbs. 2007. Binding of proprotein convertase subtilisin/kexin type 9 to epidermal growth factor-like repeat A of low density lipoprotein receptor decreases receptor recycling and increases degradation. J. Biol. Chem. 282: 18602–12.

3. Maxwell, K. N., E. A. Fisher, and J. L. Breslow. 2005. Overexpression of PCSK9 accelerates the degradation of the LDLR in a post-endoplasmic reticulum compartment. Proc. Natl. Acad. Sci. 102: 2069–2074.

4. Park, S. W., Y.-A. Moon, and J. D. Horton. 2004. Post-transcriptional regulation of low density lipoprotein receptor protein by proprotein convertase subtilisin/kexin type 9a in mouse liver. J. Biol. Chem. 279: 50630–8.

5. Goldstein, J. L., and M. S. Brown. 2015. A Century of Cholesterol and Coronaries: From Plaques to Genes to Statins. Cell. 161: 161–172.

6. Zhao, Z., Y. Tuakli-Wosornu, T. A. Lagace, L. Kinch, N. V Grishin, J. D. Horton, J. C. Cohen, and H. H. Hobbs. 2006. Molecular characterization of loss-of-function mutations in PCSK9 and identification of a compound heterozygote. Am. J. Hum. Genet. 79: 514–23.

7. Hooper, A. J., A. D. Marais, D. M. Tanyanyiwa, and J. R. Burnett. 2007. The C679X mutation in PCSK9 is present and lowers blood cholesterol in a Southern African population. Atherosclerosis. 193: 445–8.

8. Stein, E. A., S. Mellis, G. D. Yancopoulos, N. Stahl, D. Logan, W. B. Smith, E. Lisbon, M. Gutierrez, C. Webb, R. Wu, Y. Du, T. Kranz, E. Gasparino, and G. D. Swergold. 2012. Effect of a monoclonal antibody to PCSK9 on LDL cholesterol. N. Engl. J. Med. 366: 1108–18.

9. Sabatine, M. S., R. P. Giugliano, S. D. Wiviott, F. J. Raal, D. J. Blom, J. Robinson, C. M. Ballantyne, R. Somaratne, J. Legg, S. M. Wasserman, R. Scott, M. J. Koren, and E. A. Stein. 2015. Efficacy and Safety of Evolocumab in Reducing Lipids and Cardiovascular Events. N. Engl. J. Med. 372: 1500–1509.

10. Sabatine, M. S., R. P. Giugliano, A. C. Keech, N. Honarpour, S. D. Wiviott, S. A. Murphy, J. F. Kuder, H. Wang, T. Liu, S. M. Wasserman, P. S. Sever, T. R. Pedersen, and FOURIER Steering Committee and Investigators. 2017. Evolocumab and Clinical Outcomes in Patients with Cardiovascular Disease. N. Engl. J. Med. 376: 1713–1722.

11. Fonarow, G. C., A. C. Keech, T. R. Pedersen, R. P. Giugliano, P. S. Sever, P. Lindgren, B. van Hout, G. Villa, Y. Qian, R. Somaratne, and M. S. Sabatine. 2017. Cost-effectiveness of Evolocumab Therapy for Reducing Cardiovascular Events in Patients With Atherosclerotic Cardiovascular Disease. JAMA Cardiol. 2: 1069–78.

12. Kazi, D. S., J. Penko, P. G. Coxson, A. E. Moran, D. A. Ollendorf, J. A. Tice, and K. Bibbins-Domingo. 2017. Updated Cost-effectiveness Analysis of PCSK9 Inhibitors Based on the Results of the FOURIER Trial. JAMA. 318: 748–50.

13. Mullard, A. 2017. Nine paths to PCSK9 inhibition. Nat. Rev. Drug Discov. 16: 299–301.

14. Seidah, N. G., M. Chrétien, and M. Mbikay. 2018. The ever-expanding saga of the proprotein convertases and their roles in body homeostasis. Curr. Opin. Lipidol. 29: 144–150.

15. Seidah, N. G., M. Abifadel, S. Prost, C. Boileau, and A. Prat. 2017. The Proprotein Convertases in Hypercholesterolemia and Cardiovascular Diseases: Emphasis on Proprotein Convertase Subtilisin/Kexin 9. Pharmacol. Rev. 69: 33–52.

16. Sun, H., A. Samarghandi, N. Zhang, Z. Yao, M. Xiong, and B.-B. Teng. 2012. Proprotein convertase subtilisin/kexin type 9 interacts with apolipoprotein B and prevents its intracellular degradation, irrespective of the low-density lipoprotein receptor. Arterioscler. Thromb. Vasc. Biol. 32: 1585–95.

17. Kosenko, T., M. Golder, G. Leblond, W. Weng, and T. A. Lagace. 2013. Low density lipoprotein binds to proprotein convertase subtilisin/kexin type-9 (PCSK9) in human plasma and inhibits PCSK9-mediated low density lipoprotein receptor degradation. J. Biol. Chem. 288: 8279–88.

18. Kwon, H. J., T. A. Lagace, M. C. McNutt, J. D. Horton, and J. Deisenhofer. 2008. Molecular basis for LDL receptor recognition by PCSK9. Proc. Natl. Acad. Sci. U. S. A. 105: 1820–1825.

19. Benjannet, S., Y. G. L. Saavedra, J. Hamelin, M.-C. Asselin, R. Essalmani, A. Pasquato, P. Lemaire, G. Duke, B. Miao, F. Duclos, R. Parker, G. Mayer, and N. G. Seidah. 2010. Effects of the prosegment and pH on the activity of PCSK9: evidence for additional processing events. J. Biol. Chem. 285: 40965–78.

20. Dubuc, G., A. Chamberland, H. Wassef, J. Davignon, N. G. Seidah, L. Bernier, and A. Prat. 2004. Statins Upregulate PCSK9, the Gene Encoding the Proprotein Convertase Neural Apoptosis-Regulated Convertase-1 Implicated in Familial Hypercholesterolemia. Arterioscler. Thromb. Vasc. Biol. 24: 1454–9.

21. Jeong, H. J., H.-S. Lee, K.-S. Kim, Y.-K. Kim, D. Yoon, and S. W. Park. 2008. Sterol-dependent regulation of proprotein convertase subtilisin/kexin type 9 expression by sterol-regulatory element binding protein-2. J. Lipid Res. 49: 399–409.

22. Shende, V. R., M. Wu, A. B. Singh, B. Dong, C. F. K. Kan, and J. Liu. 2015. Reduction of circulating PCSK9 and LDL-C levels by liver-specific knockdown of HNF1α in normolipidemic mice. J. Lipid Res. 56: 801–809.

23. Tao, R., X. Xiong, R. A. DePinho, C.-X. Deng, and X. C. Dong. 2013. FoxO3 Transcription Factor and Sirt6 Deacetylase Regulate Low Density Lipoprotein (LDL)-cholesterol Homeostasis via Control of the Proprotein Convertase Subtilisin/Kexin Type 9 (*Pcsk9*) Gene Expression. J. Biol. Chem. 288: 29252–29259.

24. Chen, X.-W., H. Wang, K. Bajaj, P. Zhang, Z.-X. Meng, D. Ma, Y. Bai, H.-H. Liu, E. Adams, A. Baines, G. Yu, M. A. Sartor, B. Zhang, Z. Yi, J. Lin, S. G. Young, R. Schekman, and D. Ginsburg. 2013. SEC24A deficiency lowers plasma cholesterol through reduced PCSK9 secretion. Elife. 2: e00444.

25. Seidah, N. G., S. Benjannet, L. Wickham, J. Marcinkiewicz, S. B. Jasmin, S. Stifani, A. Basak, A. Prat, and M. Chretien. 2003. The secretory proprotein convertase neural apoptosis-regulated convertase 1 (NARC-1): liver regeneration and neuronal differentiation. Proc. Natl. Acad. Sci. U. S. A. 100: 928–33.

26. Benjannet, S., D. Rhainds, R. Essalmani, J. Mayne, L. Wickham, W. Jin, M.-C. Asselin, J. Hamelin, M. Varret, D. Allard, M. Trillard, M. Abifadel, A. Tebon, A. D. Attie, D. J. Rader, C. Boileau, L. Brissette, M. Chrétien, A. Prat, and N. G. Seidah. 2004. NARC-1/PCSK9 and its natural mutants: zymogen cleavage and effects on the low density lipoprotein (LDL) receptor and LDL cholesterol. J. Biol. Chem. 279: 48865–75.

27. Chorba, J. S., and K. M. Shokat. 2014. The proprotein convertase subtilisin/kexin type 9 (PCSK9) active site and cleavage sequence differentially regulate protein secretion from proteolysis. J. Biol. Chem. 289: 29030–43.

28. Chorba, J. S., A. M. Galvan, and K. M. Shokat. 2017. Stepwise processing analyses of the single-turnover PCSK9 protease reveal its substrate sequence specificity and link clinical genotype to lipid phenotype. J. Biol. Chem. 293: 1875–1886.

29. Maxwell, K. N., and J. L. Breslow. 2004. Adenoviral-mediated expression of Pcsk9 in mice results in a low-density lipoprotein receptor knockout phenotype. Proc. Natl. Acad. Sci. U. S. A. 101: 7100–7105.

30. McNutt, M. C., H. J. Kwon, C. Chen, J. R. Chen, J. D. Horton, and T. a Lagace. 2009. Antagonism of secreted PCSK9 increases low density lipoprotein receptor expression in HepG2 cells. J. Biol. Chem. 284: 10561–70.

31. Gustafsen, C., D. Olsen, J. Vilstrup, S. Lund, A. Reinhardt, N. Wellner, T. Larsen, C. B. F. Andersen, K. Weyer, J. Li, P. H. Seeberger, S. Thirup, P. Madsen, and S. Glerup. 2017. Heparan sulfate proteoglycans present PCSK9 to the LDL receptor. Nat. Commun. 8: 503.

32. Gibson, D. G., L. Young, R.-Y. Chuang, J. C. Venter, C. A. Hutchison, and H. O. Smith. 2009. Enzymatic assembly of DNA molecules up to several hundred kilobases. Nat. Methods. 6: 343–5.

33. Liu, H., and J. H. Naismith. 2008. An efficient one-step site-directed deletion, insertion, single and multiple-site plasmid mutagenesis protocol. BMC Biotechnol. 8: 91.

34. Weisgraber, K. H., T. L. Innerarity, and R. W. Mahley. 1978. Role of lysine residues of plasma lipoproteins in high affinity binding to cell surface receptors on human fibroblasts. J. Biol. Chem. 253: 9053–62.

35. Mahley, R. W., K. H. Weisgraber, G. W. Melchior, T. L. Innerarity, and K. S. Holcombe. 1980. Inhibition of receptor-mediated clearance of lysine and arginine-modified lipoproteins from the plasma of rats and monkeys. Proc. Natl. Acad. Sci. U. S. A. 77: 225–9.

36. Graham, J. M., B. A. Griffin, I. G. Davies, and J. A. Higgins. 2001. Methods Mol Med. 52: 51–9.

37. Min, D. K., H. S. Lee, N. Lee, C. J. Lee, H. J. Song, G. E. Yang, D. Yoon, and S. W. Park. 2015. In silico screening of chemical libraries to develop inhibitors that hamper the interaction of PCSK9 with the LDL receptor. Yonsei Med. J. 56: 1251–1257.

38. Lagace, T. A., D. E. Curtis, R. Garuti, M. C. McNutt, S. W. Park, H. B. Prather, N. N. Anderson, Y. K. Ho, R. E. Hammer, and J. D. Horton. 2006. Secreted PCSK9 decreases the number of LDL receptors in hepatocytes and in livers of parabiotic mice. J. Clin. Invest. 116: 2995–3005.

39. Steinman, R. M., I. S. Mellman, W. A. Muller, and Z. A. Cohn. 1983. Endocytosis and the recycling of plasma membrane. J. Cell Biol. 96: 1–27.

40. Goldstein, J. L., S. K. Basu, G. Y. Brunschede, and M. S. Brown. 1976. Release of low density lipoprotein from its cell surface receptor by sulfated glycosaminoglycans. Cell. 7: 85–95.

41. Vijayagopal, P., S. R. Srinivasan, B. Radhakrishnamurthy, and G. S. Berenson. 1981. Interaction of serum lipoproteins and a proteoglycan from bovine aorta. J. Biol. Chem. 256: 8234–41.

42. Luna Saavedra, Y. G., J. Zhang, and N. G. Seidah. 2013. PCSK9 Prosegment Chimera as Novel Inhibitors of LDLR Degradation. PLoS One. 8: e72113.

43. Homer, V. M., A. D. Marais, F. Charlton, A. D. Laurie, N. Hurndell, R. Scott, F. Mangili, D. R. Sullivan, P. J. Barter, K.-A. Rye, P. M. George, and G. Lambert. 2008. Identification and characterization of two non-secreted PCSK9 mutants associated with familial hypercholesterolemia in cohorts from New Zealand and South Africa. Atherosclerosis. 196: 659–666.

44. Berneis, K. K., and R. M. Krauss. 2002. Metabolic origins and clinical significance of LDL heterogeneity. J. Lipid Res. 43: 1363–79.

45. Tabas, I., K. J. Williams, and J. Boren. 2007. Subendothelial Lipoprotein Retention as the Initiating Process in Atherosclerosis: Update and Therapeutic Implications. Circulation. 116: 1832–1844.

46. Otvos, J. D., S. Mora, I. Shalaurova, P. Greenland, R. H. Mackey, and D. C. Goff. 2011. Clinical implications of discordance between low-density lipoprotein cholesterol and particle number. J. Clin. Lipidol. 5: 105–113.

47. Austin, M. A., J. L. Breslow, C. H. Hennekens, J. E. Buring, W. C. Willett, and R. M. Krauss. 1988. Low-density lipoprotein subclass patterns and risk of myocardial infarction. JAMA. 260: 1917–21.

